# Environmental gradients decouple demographic and adaptive connectivity in a highly mobile coastal marine species

**DOI:** 10.1101/2025.07.15.665014

**Authors:** Chris Brauer, Andrea Bertram, Jonathan Sandoval-Castillo, Anthony Fowler, Justin Bell, Paul Hamer, Maren Wellenreuther, Luciano B. Beheregaray

**Affiliations:** Molecular Ecology Laboratory, College of Science and Engineering, Flinders University, Bedford Park, SA, Australia; Aquatic Sciences, South Australian Research and Development Institute, Henley Beach, SA, Australia; Victorian Fisheries Authority, Queenscliff, Vic, Australia; Pacific Community, Noumea, New Caledonia; The New Zealand Institute for Plant and Food Research Limited, Nelson, New Zealand; The School of Biological Sciences, University of Auckland, Auckland, New Zealand

**Keywords:** ecological genomics, marine biogeography, teleost, recruitment dynamics, fisheries management

## Abstract

Understanding how eco-evolutionary processes shape genetic variation and persistence in marine species with highly variable recruitment dynamics and dispersal potential remains a fundamental challenge, particularly when considering the interplay between gene flow and local adaptation. Here, we employed a seascape genomics approach to investigate population connectivity and local adaptation in Australasian snapper (*Chrysophrys auratus*, Sparidae) along 1500 km of the environmentally heterogeneous southern Australian coastline. Using 14,699 SNPs, we identified two distinct regional populations aligned with known biogeographical regions. Genotype-environment association analyses revealed 855 candidate adaptive loci associated with environmental variation, including temperature, salinity, and primary productivity. Connectivity analyses using neutral markers indicated high gene flow throughout both eastern and western regions, while candidate adaptive loci revealed substantially reduced connectivity, especially for Northern Spencer Gulf and West Coast populations. This pattern of locally restricted gene flow at adaptive genomic regions suggests that strong environmental gradients are driving adaptive divergence despite high overall connectivity. Our results support contingent migration as a potential mechanism modulating the balance between local adaptation and gene flow in this economically and ecologically important marine species. These findings might also have implications for regional management of other coastal fisheries that are experiencing substantial declines. As climate change alters coastal marine environments around the world, the dynamics of local recruitment, site fidelity and local adaptation are expected to change. This highlights the importance of integrating knowledge about eco-evolutionary processes into marine resource management, fisheries stock assessment, and restocking and stock enhancement activities.

## Introduction

Understanding how evolutionary processes such as gene flow and local adaptation shape the distribution of marine populations are enduring challenges in evolutionary biology and for fisheries management (Bernatchez et al., 2017; Grummer et al., 2019). This is particularly true for many marine species characterised by large population sizes, high fecundity and high connectivity (Gagnaire et al., 2015), including those where recruitment dynamics is influenced by sweepstakes reproductive success (Árnason et al., 2024). In these cases, extensive gene flow could limit local adaptation, even across vast geographical ranges (Lenormand, 2002; Nielsen, HemmerLHansen, Larsen, & Bekkevold, 2009). Despite these challenges, growing evidence indicates that locally adapted genetic variation can be maintained in species with high gene flow (Han et al., 2020).

Marine fishes provide ideal natural systems to examine the genomic basis of local adaptation with gene flow, particularly in the context of fisheries resources. The management consequences of local adaptation for widespread fisheries stocks are, however, relatively unexplored (Andersson et al., 2024; Grummer et al., 2019). Highly dispersive marine species often span extremely heterogeneous environments, yet conventional stock assessments may fail to capture this complexity. Additionally, fisheries management and stock assessments are often constrained by government jurisdictional boundaries that reflect political or administrative divisions rather than biologically- or ecologically-relevant population structure (Andersson et al., 2024). This potential disparity between management units and biological populations can lead to localized overfishing, inadequate protection of important spawning or nursery areas, or failure to account for source–sink dynamics occurring at larger spatial scales (Berger et al., 2021).

Seascape genomics offers a powerful approach to disentangle demographic and ecological evolutionary processes in these systems (Gagnaire et al., 2015). This interdisciplinary field can clarify ecological and evolutionary factors shaping population structure and connectivity of widespread fisheries species (Xuereb, d’Aloia, Andrello, Bernatchez, & Fortin, 2021). By examining how the environment influences genetic variation and gene flow, seascape genomics can reveal subtle population structure that may be overlooked by previous genetic methods or stock assessment approaches (SandovalLCastillo, Robinson, Hart, Strain, & Beheregaray, 2018). Atlantic herring (*Clupea harengus*) provide a clear example where, despite limited evidence for population structure at neutral loci, seascape genomics revealed fine-scale differentiation associated with ecological adaptation that helped to resolve mismatches between biological populations and fisheries stock management (Han et al., 2020). Similarly, temperature-associated adaptive variation was identified in the commercially important yellow croaker (*Larimichthys crocea*) along the Chinese coast, challenging the existing stock assessment and highlighting the need for more nuanced management strategies (Chen et al., 2023).

Coastal marine species often exhibit limited population structure across continental-scale distributions even if they encompass a wide range of environments. If patterns of dispersal are locally restricted or temporally variable within these large populations, environmental heterogeneity can, however, create opportunities for natural selection to drive local adaptation. For these reasons, spatially and temporally heterogeneous coastal habitats, such as the zonal coastal boundary of southern Australia, provide ideal opportunities to test for the role of ecologically divergent natural selection in driving adaptation in marine species (Ruzzante et al., 2006; SandovalLCastillo & Beheregaray, 2020).

Here we examine the roles of regional and fine-scale connectivity linked to variable recruitment dynamics in shaping population connectivity and adaptation in the Australasian snapper (*Chrysophrys auratus*), a highly mobile coastal marine species. The focal region of our study spans a range of heterogeneous embayment and open coastal environments along more than 1500 km of the southern Australian coast (Ridgway & Condie, 2004). Snapper plays a pivotal role in the commercial and recreational fisheries sector of southern Australia, as well as holding considerable ecological and cultural value across its Indo-Pacific distribution (Parsons et al., 2014). Recent declines in the productivity and status of some snapper stocks have prompted significant shifts in management practices, including an ongoing moratorium, now in its sixth year, on commercial and recreational fishing in much of South Australia (Drew et al., 2022; Fowler et al., 2020). To inform and assess management of the species it is critically important to understand how connectivity and environmental variation shape snapper stocks subject to both fishing pressure and climate change. Several genetic studies have revealed evidence for fine-scale population structure and provide a basis for the hypothesis that selection, imposed by steep environmental gradients, could counteract high gene flow and lead to local adaptation of snapper populations (Bernal-Ramírez, Adcock, Hauser, Carvalho, & Smith, 2003; Bertram et al., 2023; Gardner, Chaplin, Potter, Fairclough, & Jackson, 2017). Spawning aggregations and nursery areas across southern Australia concentrate in three major embayments: northern Spencer Gulf, northern Gulf St. Vincent, and Port Phillip Bay. Previous genetic analyses suggest these populations exhibit local recruitment and site fidelity (Bertram et al., 2023), traits that favour the evolution of local adaptation. This is supported by early molecular evidence for local adaptation in other embayment populations, including Shark Bay, Western Australia (Johnson, Creagh, & Moran, 1986), and New Zealand (Smith, 1979; Smith & Francis, 1983). Adaptive traits such as larval survival, growth and metabolism have been strongly linked to temperature and salinity in snapper (Fielder, Bardsley, Allan, & Pankhurst, 2005; McMahon, Parsons, Donelson, Pether, & Munday, 2020; Wellenreuther, Le Luyer, Cook, Ritchie, & Bernatchez, 2019), and genomic variation is known to underpin these traits (Ashton, Hilario, Jaksons, Ritchie, & Wellenreuther, 2019a; Ashton, Ritchie, & Wellenreuther, 2019b; Sandoval-Castillo, Beheregaray, & Wellenreuther, 2022).

In this study, we predicted that genomic variation should not only reflect demographic factors such as large population sizes and high gene flow, but also natural selection in response to environmental heterogeneity. To test these predictions, we built on recent evaluations of snapper stock structure and population size estimates (Bertram et al., 2022; Bertram et al., 2023; Bertram et al., 2024) and examined how environmental parameters may influence patterns of genomic variation and connectivity in snapper populations along the southern Australian coast. Considering the social, economic, and commercial importance of southern Australian snapper stocks, we also aimed to assess the findings in the context of informing and refining regional fisheries management strategies which are fundamental for sustainable harvesting and species persistence.

## Methods

### Southern Australian coastal environment

The focal region of this study, southern Australia, comprises a wide range of coastal and near-shore marine habitats and includes three major embayments. Spencer Gulf and the adjacent Gulf St. Vincent are large inverse estuaries in South Australia, and Port Phillip Bay is a shallow, enclosed bay just south of Melbourne, Victoria (Figure 1). Broadly, environmental conditions within these embayments differ greatly from open coastal habitats, with cooler minimum sea surface temperature (SST), warmer maximum SST and reduced current velocities. A west-to-east temperature gradient characterises the region, with the warmest waters along the west coast and coolest in the eastern embayments. Regional variation in climate, topography and oceanography create unique local environments within each embayment. Climate in the east is wetter and cooler than the more arid South Australia. As a result, Port Phillip Bay and Western Port Bay exhibit the coolest SST and receive greater freshwater inputs from rivers, leading to the highest primary productivity in the region and consistently low salinity. In South Australia, Spencer Gulf is characterised by extreme salinity gradients, particularly in the north where values exceed 45 ppt in summer, combined with large seasonal variation in SST (∼12–24°C) (Nunes & Lennon, 1986). Gulf St. Vincent experiences intermediate conditions that are less saline than Spencer Gulf but saltier than coastal waters. Coastal current circulation outside the Gulfs is dominated by the Leeuwin current system that transports tropical water poleward along the west coast of Australia and eastward along the southern continental shelf, before joining the South Australian Current, and eventually heading south along Tasmania’s west coast as the Zeehan Current (Richardson, Middleton, Kyser, James, & Opdyke, 2019; Ridgway & Condie, 2004). The strength and impact of this current system is highly variable and shapes seasonal patterns of temperature, salinity and primary productivity across the region. Stronger currents during winter transport warm, low-salinity water to the east, while reduced Leeuwin current flows and coastal upwellings incorporate cooler, more saline and nutrient-rich water during summer (Richardson et al., 2019; Ridgway & Condie, 2004; Shute et al., 2022).

**Figure 1.**
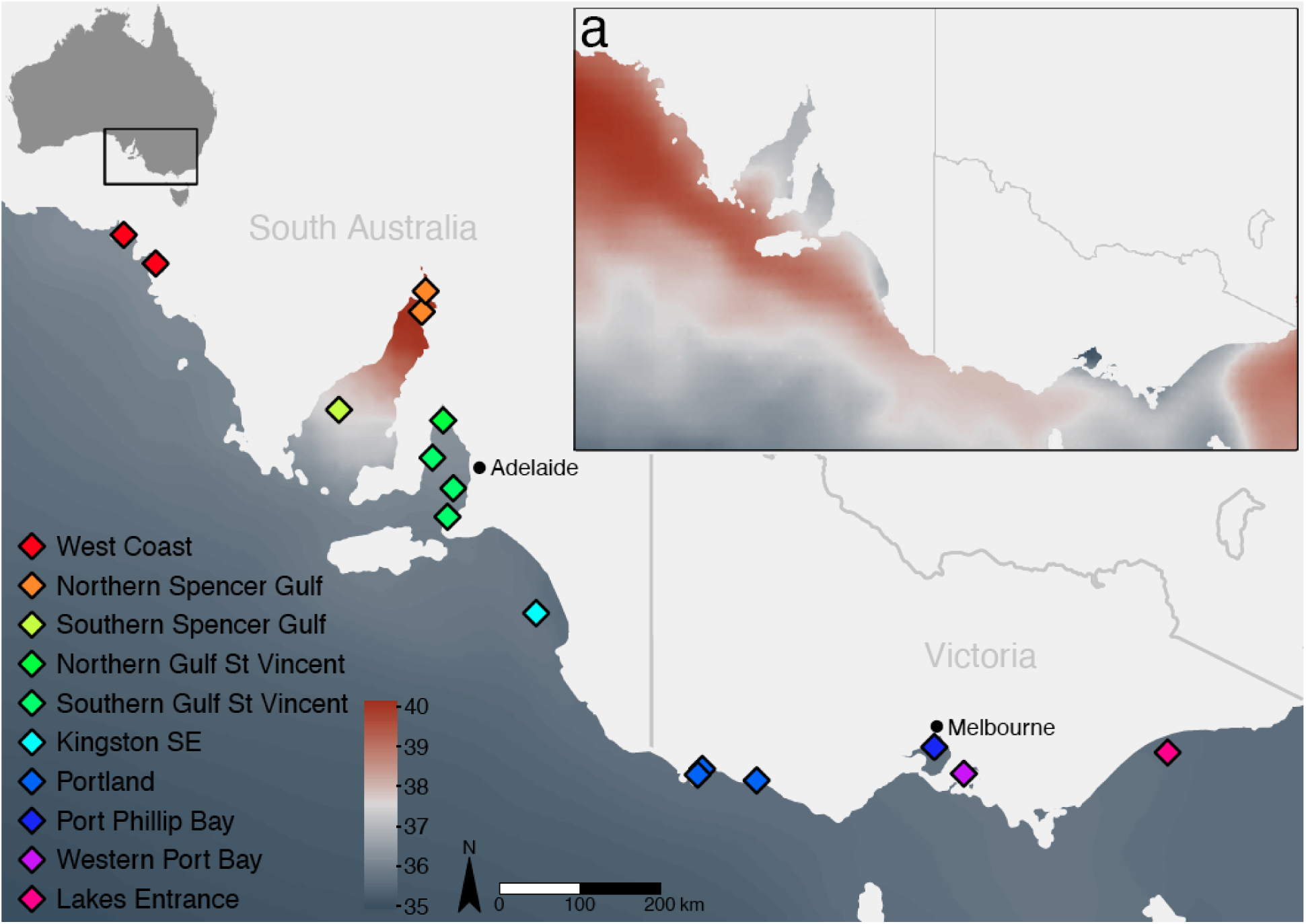
Sampling sites across southern Australia and spatial heterogeneity in mean sea surface salinity (main panel), and (a) minimum sea surface temperature.

### Sampling and genomic data collection

Our sampling design builds on previous studies that clarified neutral population genetic structure and stock boundaries of snapper along the western (Bertram et al., 2022) and the southeastern (Bertram et al., 2023; Bertram et al., 2024) coasts of Australia. A total of 448 snapper (*Chrysophrys auratus*) collected from 23 individual sampling sites along the southern Australian coast were used. These sites were grouped into ten regional locations based on proximity and similarity of environmental profiles. To ensure adequate sample sizes for allele-frequency estimation, individual sites containing fewer than six samples were merged with the nearest neighbour. This resulted in 16 pooled sites representing the ten regional locations (Figure 1; Figure S1; Table S1). All the research complies with applicable laws on sampling from natural populations. We extracted DNA from fin clips using a salting-out method (Sunnucks & Hales 1996) and used double-digest restriction site-associated DNA (ddRAD) sequencing to generate genomic data. We generated ddRAD libraries of 96 multiplexed samples, following the protocol outlined in Peterson et al. (2012) with modifications described in our previous work (Brauer, Hammer, & Beheregaray, 2016).

Paired-end 150bp sequencing was conducted over 15 lanes (this included samples not related to this study) of an Illumina HiSeq 4000 at Novogene (Hong Kong). Raw sequence data quality was assessed using FastQC (Andrews, 2010). Reads were demultiplexed with the process_radtags module from STACKS 2.0 (Catchen, Hohenlohe, Bassham, Amores, & Cresko, 2013) and trimmed to remove low-quality bases and adapters using TRIMMOMATIC (Bolger, Lohse, & Usadel, 2014). Reads were then mapped to a chromosome-level genome assembly (Catanach et al., 2019) using BOWTIE 2 (Langmead & Salzberg, 2012), and SNPs were called using BCFTOOLS (Narasimhan et al., 2016). We filtered the resulting SNP genotypes for quality, missing data per individual, and retained one SNP per 500bp. We also applied further filtering to generate putatively neutral and candidate adaptive datasets. Candidate adaptive loci identified by the genotype–environment association analysis (described below) were separated, and the remaining putatively neutral SNPs were filtered for departure from Hardy–Weinberg equilibrium (FDR of 0.05) using the gl.filter.hwe function in the *dartR* package. Filtering parameters and the number of SNPs retained following each step are described in Table S2.

### Genetic diversity, population structure and connectivity

We estimated population genetic diversity parameters, including the number of alleles, percentage of polymorphic loci, heterozygosity, and inbreeding coefficient, using the *hierfstat* R package (Goudet, 2005). We examined patterns of neutral population structure and admixture using using *hierfstat* to estimate pairwise *F*_ST_ and ADMIXTURE (Alexander, Novembre, & Lange, 2009). An analysis of molecular variance (AMOVA) was performed to further assess hierarchical population structure among major sampled regions, among sites within regions, among individuals within sites, and within individuals. The poppr.amova function in the *poppr* R package (Kamvar, Tabima, & Grünwald, 2014) was used for the molecular variance estimates and significance values were estimated using the *ade4* randtest function (Dray & Dufour, 2007) with 1000 permutations. We quantified gene flow among sites within each of the eastern and western regional populations with the directional relative-migration (Nm) approach of Sundqvist (2016), as implemented by the divMigrate function in the *diveRsity* R package (Keenan, McGinnity, Cross, Crozier, & Prodöhl, 2013). This approach first calculates pairwise differentiation (*G*_ST_), then converts these values into unit-less estimates of relative migrants per generation and assigns directionality by contrasting each population’s allele frequencies with those of a hypothetical mixed migrant pool.

Analyses were run separately for the neutral and putatively adaptive SNP datasets, and statistical support was evaluated with 1,000 bootstrap resamples.

### Local Adaptation

Environmental heterogeneity across the study area was summarised with 31 raster layers at 5-arc-min resolution (∼9 km at the equator) obtained from BioORACLE v2.2 (Assis et al., 2018; Tyberghein et al., 2012), supplemented by bathymetry from BioORACLE v1.0 (Table S3). Each layer represents long-term summary statistics (e.g. mean, minimum, maximum, or range), calculated over a baseline period of 2000-2014 and derived either from the ECMWF ORAP5.0 ocean reanalysis (physical variables), or the PISCES biogeochemical hind-cast (nutrients and productivity). Two variables rely on different archives and time periods, pH (1910-2007 observations, World Ocean Database) and calcite (2002-2009 MODIS-Aqua).

We performed preliminary analyses using the vifcor and vifstep functions in the *usdm* R package (Naimi, Hamm, Groen, Skidmore, & Toxopeus, 2014) to remove correlated (Pearson r > 0.7) and colinear (VIF > 10) variables. The retained environmental variables were finally converted to z-scores to standardize across different measurement scales and ensure comparable contributions to downstream analyses.

To detect a genetic signal of local adaptation, we performed genotype-environment association analyses using redundancy analysis (RDA) implemented with the *rda* function in the vegan R package (Oksanen et al., 2018). To examine relationships between SNP allele frequencies and environmental data, we ran an initial partial RDA where we controlled for spatial population structure using a matrix of allele frequency covariance (Ω) estimated with Baypass (Gautier, 2015). Candidate adaptive loci significantly associated with environmental variation were identified using the Mahalanobis distance statistical approach proposed by Capblancq et al. (2018). A second RDA was then performed using the candidate adaptive loci and individual SNP genotypes to visualise the distribution of genotype-environment associations within populations.

Functional annotations of candidate loci were performed using SnpEff to assess genomic position and predicted effects (Cingolani et al., 2012). We then extracted open reading frames (ORFs) from 600 bp flanking sequences around each SNP and searched these against the SwissProt teleost database using DIAMOND BLASTx (Buchfink, Xie, & Huson, 2015), applying an e-value threshold of 1 × 10L¹L and retaining up to five hits per query. Top-scoring matches were queried through the UniProt REST API to retrieve Gene Ontology (GO) terms and KEGG pathways, retaining only those with experimental or curated evidence codes (EXP, IDA, IPI, IMP, IGI, IEP, TAS, IC) or high-confidence computational inference (ISS, IEA). To identify enriched functional categories, we performed over-representation analysis using the *clusterprofiler* R package (Yu, Wang, Han, & He, 2012). Annotated candidate loci were compared against all annotated loci using hypergeometric tests, with false discovery rate controlled by the Benjamini–Hochberg procedure (FDR < 0.1). Categories containing fewer than five or more than 500 genes were excluded to avoid unstable or overly general terms. The genomic distribution of candidate loci and enriched functional categories was visualized using the ggmanh R package (Lee, 2022).

## Results

### Sampling and genomic data collection

The Illumina HiSeq 4000 sequencing generated 5.05 billion raw sequences, and 3.36 billion (mean per sample = 7.51M, min = 0.87M, max = 21.38M) were retained. After filtering, we retained 448 individuals genotyped at 14,699 single nucleotide polymorphisms (SNPs).

Filtering for Hardy–Weinberg equilibrium (HWE), retained 14,206 SNPs, and exclusion of candidate adaptive loci resulted in a putatively neutral dataset of 13,453 SNPs.

### Genetic diversity, population structure and connectivity

Estimates of genetic diversity were based on the 13,453 neutral loci, with observed heterozygosity (*H*_o_) ranging from 0.124 in southern Gulf St. Vincent to 0.138 in southern Spencer Gulf, while expected heterozygosity (*H*_e_) varied from 0.128 to 0.133. Inbreeding coefficient (*F*_is_) values were generally low, ranging from −0.036 in southern Spencer Gulf to 0.036 in northern Gulf St. Vincent, indicating minimal inbreeding within populations. Percentages of polymorphic loci ranged from 79.1% in Kingston–South East to 87.0% in Port Phillip Bay (Table 1).

**Table 1.** Genetic diversity indices based on 13,453 neutral loci. Number of individuals, N; number of alleles, N_A_; percentage polymorphic loci, %PL; observed heterozygosity, H_O_; expected heterozygosity, H_E_; and inbreeding coefficient, F_IS_ (including 95% confidence intervals, LCI and UCI).

| Site | $N$ | $N_A$ | %poly | $H_O$ | $H_E$ | $F_{IS}$ | LCI | UCI |
| --- | --- | --- | --- | --- | --- | --- | --- | --- |
| West Coast (WC) | 38 | 1.610 | 83.5 | 0.125 | 0.129 | 0.033 | 0.028 | 0.036 |
| Northern Spencer Gulf (NSG) | 40 | 1.607 | 83.3 | 0.125 | 0.128 | 0.019 | 0.016 | 0.023 |
| Southern Spencer Gulf (SSG) | 36 | 1.625 | 84.2 | 0.138 | 0.133 | -0.036 | -0.040 | -0.032 |
| Northern Gulf St Vincent (NGSV) | 34 | 1.626 | 83.4 | 0.127 | 0.131 | 0.036 | 0.032 | 0.041 |
| Southern Gulf St Vincent (SGSV) | 39 | 1.597 | 80.7 | 0.124 | 0.128 | 0.030 | 0.027 | 0.034 |
| Kingston SE (KSE) | 70 | 1.602 | 79.1 | 0.128 | 0.130 | 0.013 | 0.009 | 0.016 |
| Portland (PLD) | 69 | 1.633 | 85.6 | 0.129 | 0.132 | 0.022 | 0.019 | 0.026 |
| Port Phillip Bay (PPB) | 45 | 1.623 | 87.0 | 0.129 | 0.131 | 0.013 | 0.008 | 0.017 |
| Western Port Bay (WPB) | 37 | 1.628 | 85.1 | 0.131 | 0.132 | 0.010 | 0.006 | 0.014 |
| Lakes Entrance (LE) | 40 | 1.601 | 81.8 | 0.129 | 0.129 | 0.000 | -0.004 | 0.005 |

ADMIXTURE results indicated that the optimal number of genetic clusters was KL=L2 (Figure 2), corresponding to an eastern and a western cluster with the genetic break situated between southern Gulf St. Vincent and Kingston–South East. Despite this main division, a few migrants (three from the eastern and two from the western cluster) are evident in the ADMIXTURE plot. Pairwise *F*_ST_ values based on the 13,453 neutral loci were generally low among sites within regions but higher between eastern and western regions (mean = 0.009, range = 0-0.018). Based on the candidate adaptive loci, *F*_ST_ estimates were elevated, both among sites within each region, and among sites across regions (mean = 0.013, range = 0-0.25; Figure S2). The AMOVA analyses supported the ADMIXTURE results, attributing a significant proportion of genetic variation to differences between regions (2.1%, ΦL=L0.021, PL<L0.01), while variation among sites within regions was minimal (0.1%, ΦL=L0.001, PL<L0.001). The AMOVA based on candidate adaptive loci assigned a higher proportion of variation between sites within regions (0.4%, ΦL=L0.004, PL<L0.001), and among regions (5.9%, ΦL=L0.059, PL<L0.01), indicating stronger differentiation at loci potentially under selection (Table 2).

**Figure 2.**
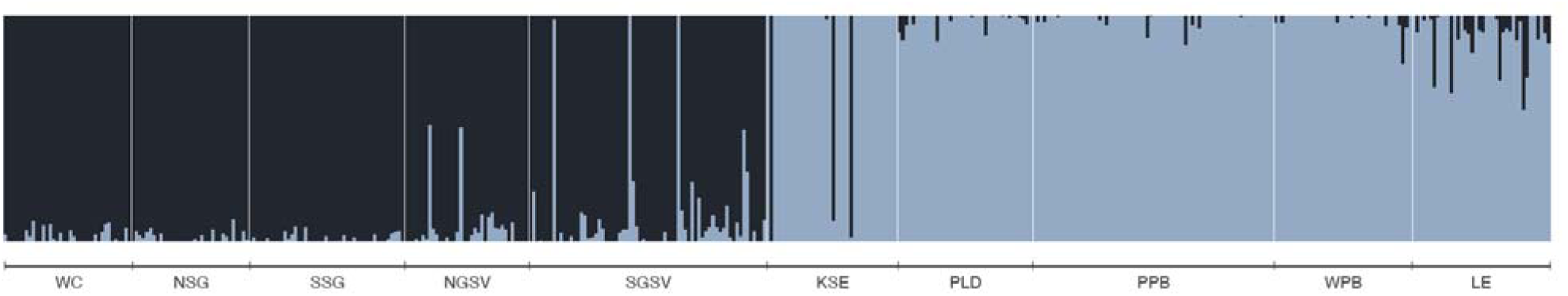
ADMIXTURE results for K=2, indicating distinct western (dark) and eastern (light) genetic clusters. Site codes refer to the locations listed in Table 1.

**Table 2.**
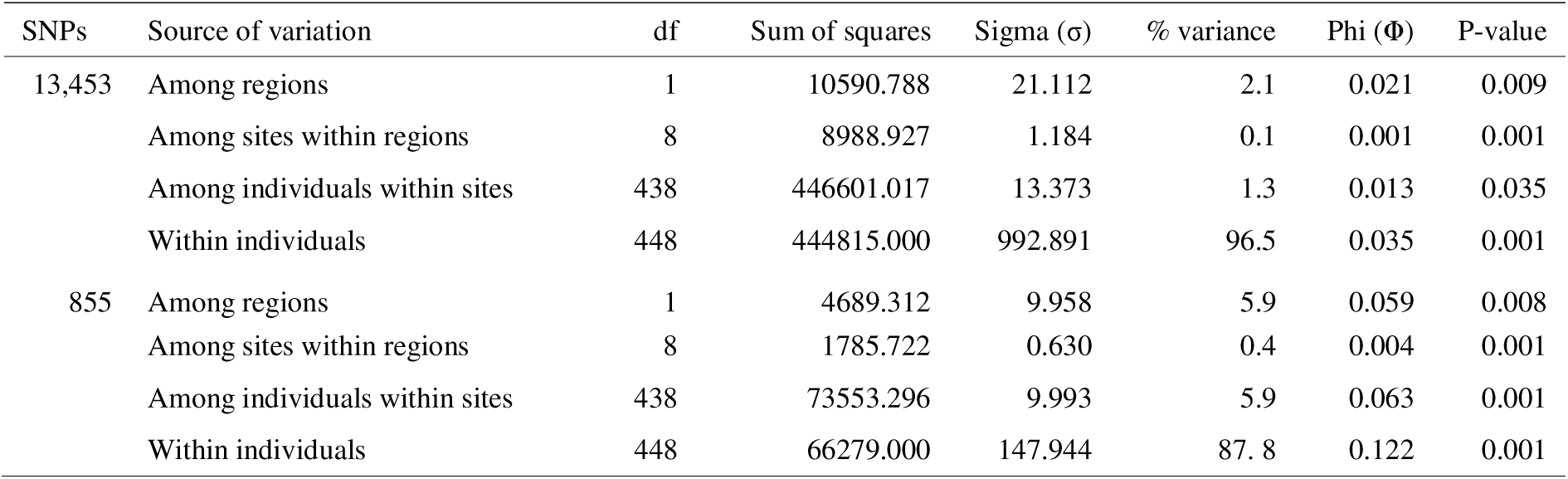
Hierarchical analysis of molecular variance (AMOVA) based on 13,453 neutral and 855 candidate loci, illustrating the distribution of genetic variation among regions (eastern, western), sites within each region, and individuals within sites, and within individuals.

The relative migration (Nm) estimates based on neutral loci for both the eastern and western regions showed high connectivity among sites (eastern mean Nm = 0.79, western mean Nm = 0.77; Figure 3a,c, Table S4-5), with particularly high connectivity evident between the southern areas of the two gulfs, Spencer Gulf and Gulf St. Vincent (mean Nm = 0.97), and between Port Phillip Bay and all other eastern sites (mean Nm = 0.89). Using the candidate loci, migration estimates for eastern region were lower (mean Nm = 0.72; Figure 3b, Table S6). For the western region, relative migration estimates again suggested the southern areas of both gulfs are highly connected (mean Nm = 0.98; Figure 3d, Table S7) but indicated much lower connectivity for all pairwise estimates involving West Coast (mean Nm = 0.53) and, particularly northern Spencer Gulf (mean Nm = 0.46).

**Figure 3.**
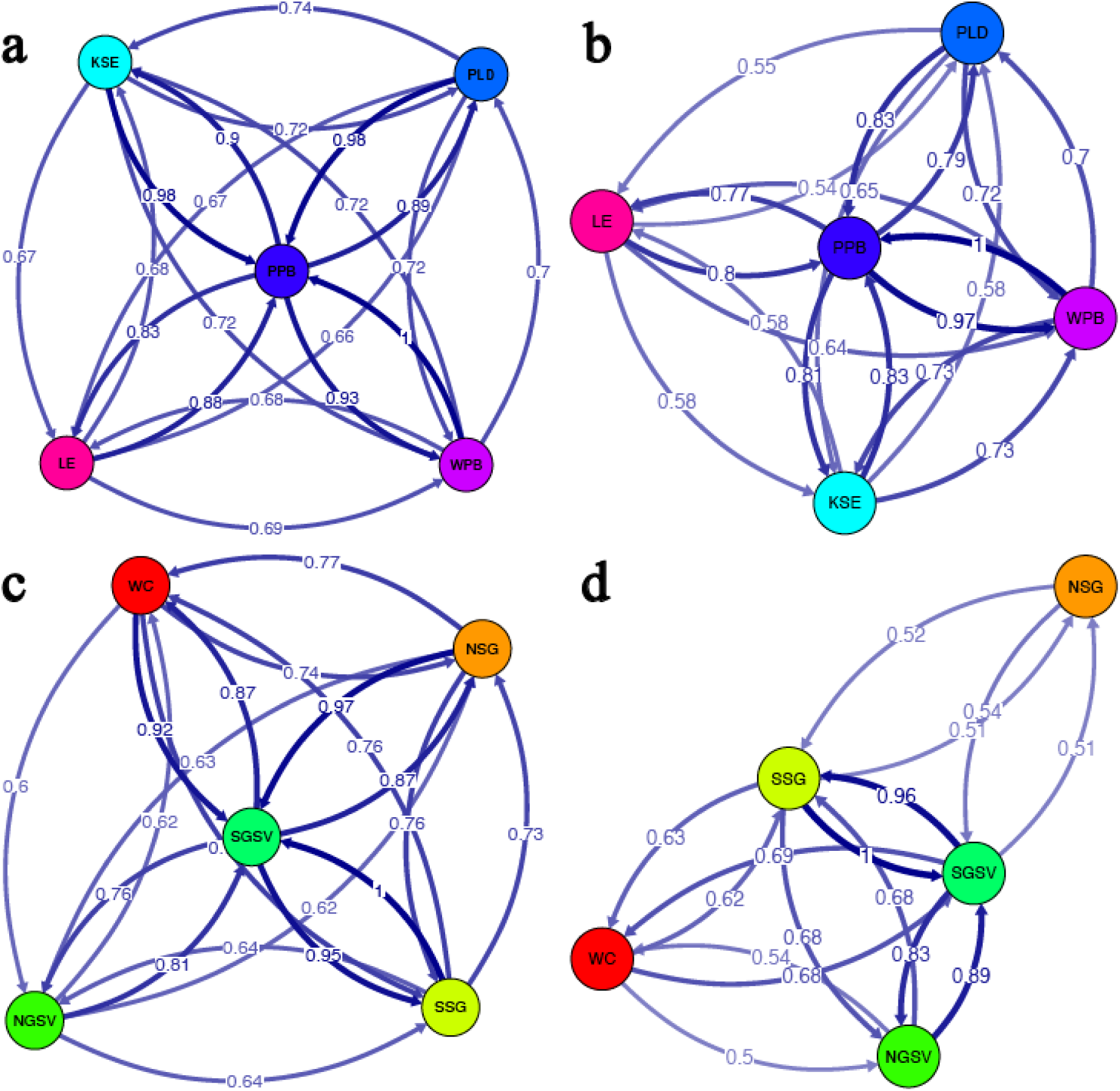
Relative migration networks for the eastern stock estimated using (a) 13,453 neutral and (b) 855 candidate adaptive SNPs, and for the western stock using (c) 13,453 neutral and (d) 855 candidate SNPs, following the method of Sundqvist (2016). Darker arrows signify higher relative migration. Estimates <0.5 are not shown.

Filtering correlated (Pearson r > 0.7) and colinear (VIF > 10) variables resulted in a final set of five environmental variables describing environmental heterogeneity across the study region (minimum sea surface temperature, mean sea surface salinity, minimum sea surface net primary productivity, pH, and calcite concentration; Figure S3, Table S8). These variables capture critical factors known to influence snapper physiology and life history, including thermal tolerance during winter months, osmoregulatory stress in hypersaline embayments, food availability during low-productivity periods, and carbonate chemistry affecting calcification processes. The partial RDA identified 855 candidate adaptive loci (5.8% of the total SNPs) significantly associated with the environment (Figure 4a). The individual-level RDA based on those 855 candidate loci (Figure 4b) revealed divergent genotype-environment associations between regions, with western populations showing strong associations with salinity and calcite (reflecting the hypersaline Spencer Gulf environment), while eastern populations were more strongly associated with temperature minima and productivity gradients.

**Figure 4.**
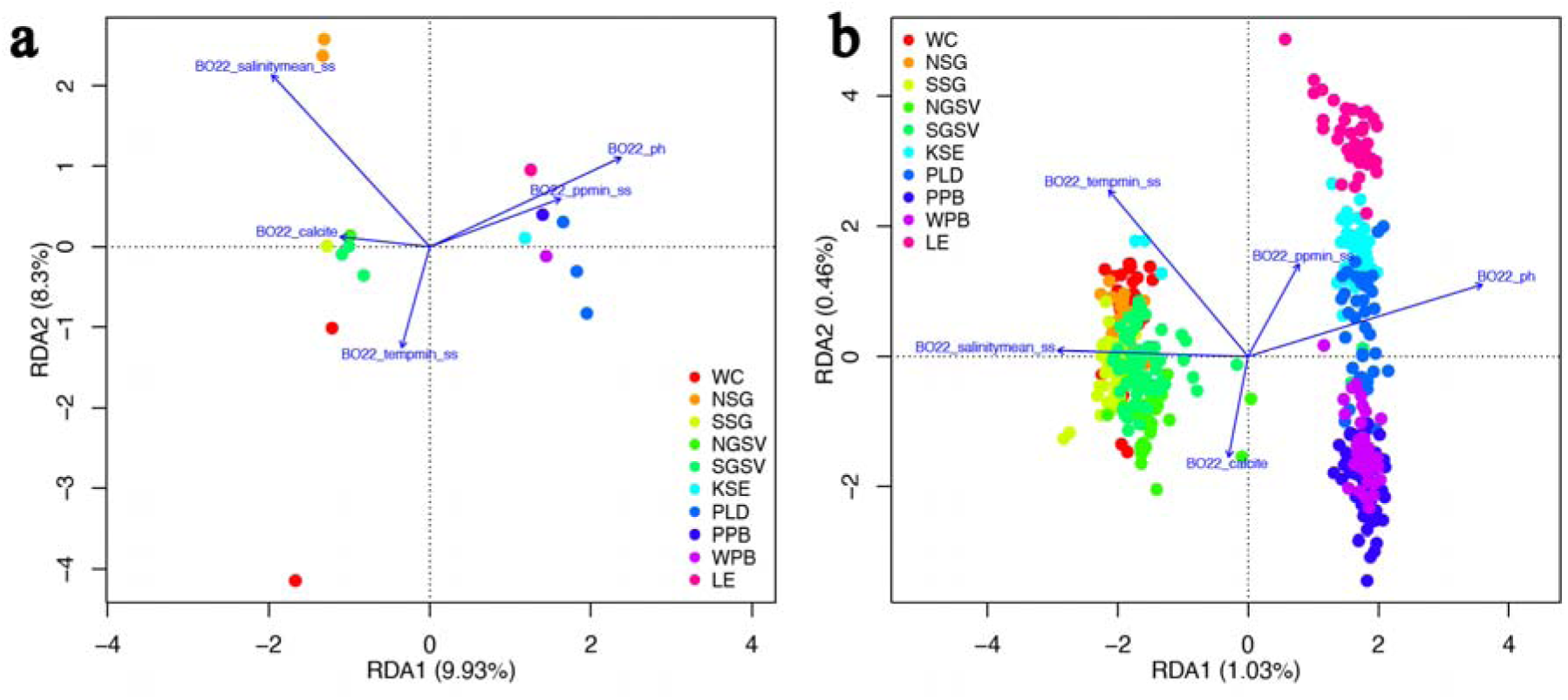
Redundancy analysis (RDA) plots summarising genotype–environment associations in Australasian snapper: (a) RDA biplot showing ordination of sampling locations based on 14,699 genome-wide SNPs after controlling for spatial structure using a population allele frequency covariance matrix. Arrows indicate the direction and strength of environmental gradients, including minimum sea surface temperature, mean sea surface salinity, minimum sea surface net primary productivity, pH, and calcite concentration; (b) Individual-level RDA biplot based on 855 putative adaptive candidate loci, illustrating the distribution of individual genotypes across sites in relation to environmental variables.

Functional annotation of these loci revealed a substantial proportion located in genic regions, with many predicted to have moderate to high impacts on gene function (Figure 5a-b). We recovered 1,188 BLAST hits covering 730 unique loci (85.4% of candidates). These matches corresponded to 562 unique proteins with 5,115 GO terms (2,640 biological process, 1,263 cellular component, and 1,212 molecular function terms) and 488 KEGG pathway annotations (Table S9). Based on these annotations, over-representation analysis identified two enriched biological process GO terms. Post-anal tail morphogenesis (GO:0036342; p = 0.0022, FDR = 0.062), was represented by three genes involved in FGF-BMP-Wnt signaling (*fgfr1a*, LG10; *tll1*, LG15; *bcl9l*, LG18), a conserved growth-factor network that regulates skeletal and muscle growth and mediates plastic responses to environmental cues such as temperature and salinity (Johnston, 2006). Hemopoiesis (GO:0030097) was also significantly enriched (p = 0.0043, FDR = 0.062), with three genes affecting red blood cell production or membrane composition (*rnf145*, LG16; *zfpm1*, LG4; *smarcal1*, LG8). No significant enrichment was detected for molecular function or cellular component categories, or for KEGG pathways. The functionally annotated candidate loci were distributed across the genome, with enriched genes found on multiple linkage groups (Figure 5c).

**Figure 5.**
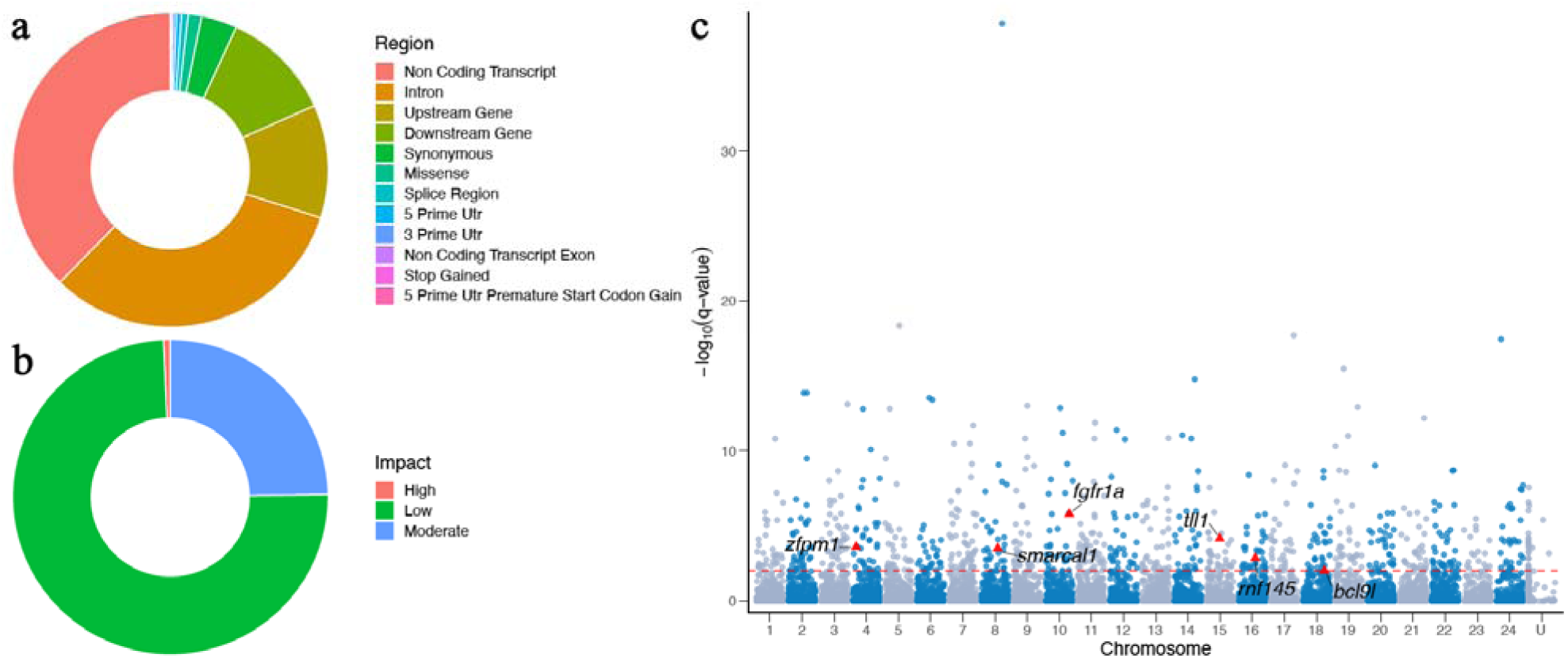
(a) Proportion of 855 candidate adaptive SNPs by genomic region; (b) Proportion of candidate SNPs in genic regions by impact on gene function; (c) Manhattan plot indicating the RDA q-values (-log_10_[*q*]) and distribution of 855 candidate adaptive SNPs associated with environmental variation (FDR 0.01; above dashed line). Red triangles highlight loci linked to six genes representing two functional gene ontology terms enriched in the candidate adaptive SNPs (FDR < 0.1), GO:0036342, post-anal tail morphogenesis (*fgfr1a*, *tll1*, *bcl9l*), and GO:0030097, hemopoiesis (*rnf145*, *zfpm1*, *smarcal1*).

## Discussion

Understanding how environmental heterogeneity shapes population structure and connectivity remains a central challenge in ecology and evolution, particularly for marine species with highly variable recruitment dynamics and dispersal potential. The relationship between gene flow and local adaptation is especially complex in marine environments where oceanographic connectivity can span vast distances while environmental gradients create strong selective pressures. Along the southern Australian coast, our findings reveal that broad environmental gradients influence patterns of dispersal and local adaptation in snapper, despite substantial gene flow among populations. We identified two distinct regional populations with minimal genetic differentiation at neutral loci among sites within each region, consistent with high demographic connectivity. However, genotype-environment association analyses identified 855 candidate adaptive loci linked to five key environmental variables that shape snapper ecology. Minimum temperature that affects spawning, growth and survival, salinity extremes challenging osmoregulation, primary productivity determining prey availability, and carbonate chemistry (pH and calcite) potentially influencing sensory systems and otolith formation (Cook, Herbert, & Jerrett, 2021; Fielder et al., 2005). A substantial proportion of these putatively adaptive SNPs are located in genic regions with potential moderate to high impacts on gene function. Functional over-representation analyses identified six candidate SNPs that mapped to two enriched GO biological process terms (post-anal tail morphogenesis; hemopoiesis), suggesting hydrodynamic niche and salinity tolerance as potential targets of selection. While relative migration estimates using neutral loci suggested high connectivity among sites within each population, using candidate loci revealed lower connectivity, particularly for West Coast and northern Spencer Gulf populations. This decoupling of demographic and adaptive connectivity highlights that substantial gene flow does not preclude local adaptation, with specific environmental stressors contributing to adaptive genetic divergence of local snapper populations.

### Strong environmental gradients influence connectivity and local adaptation

Current knowledge of snapper population dynamics across southern Australia, based on size structure, mark-recapture, and otolith studies, indicates that spatial and temporal patterns in regional connectivity mainly reflect interannual variation in recruitment to three main nursery areas: Port Phillip Bay in the east, and northern Spencer Gulf and northern Gulf St. Vincent in the west (Drew et al., 2022; Fowler et al., 2020; Hamer, Jenkins, & Gillanders, 2003). We detected evidence for occasional long-distance migration from the eastern stock into southern Gulf St. Vincent, along with more subtle, fine-scale differentiation between sub-populations within, and adjacent to the South Australian gulfs. The major genetic break we observed between the eastern and western populations aligns with the boundary between the well-described Flindersian and Maugean marine biogeographic provinces, suggesting that broad-scale climatic and oceanographic processes are important drivers of stock structure across the region (Teske, SandovalLCastillo, Waters, & Beheregaray, 2017; Waters et al., 2010). These findings are consistent with previous genetic studies (Bertram et al., 2023), and with the view that strong recruitment events drive dispersal of juvenile snapper among populations at both local and regional spatial scales (Fowler, 2016).

Although the coast of southern Australia is relatively well-studied from a biogeographic perspective (Teske et al., 2017; Waters, 2008), less is known about how its heterogeneous seascape and strong environmental gradients influence connectivity and local adaptation, particularly for teleosts. Seasonal variation in the intensity and direction of prevailing oceanographic currents can shape local environmental conditions including temperature, salinity and dissolved oxygen (Richardson et al., 2019). Such variation is also known to mediate evolutionary processes such as dispersal, recruitment, and local adaptation for a range of coastal marine species including macroalgae, sea urchins, silversides, sardines, and cetaceans (Banks et al., 2007; Barceló et al., 2022; Beheregaray & Sunnucks 2001; Coleman et al., 2011; Teske, Sandoval-Castillo, Van Sebille, Waters, & Beheregaray, 2016; Teske et al., 2021; Ward et al., 2006). For snapper, strong recruitment events can enhance demographic connectivity at both regional and local scales, from long-distance westward dispersal originating from the eastern nursery area (Fowler, 2016; Hamer & Jenkins, 2004) to more localised exchange among subpopulations within regions. However, environmental heterogeneity among nursery areas, and between embayments and adjacent coastal waters, likely imposes strong selection on migrants that may favour survival of local recruits (Rankin & Sponaugle, 2011). These patterns are consistent with a migration-selection balance, where exceptional recruitment pulses are offset by selection against maladapted immigrants, maintaining local adaptation across environmental gradients (Yeaman & Whitlock, 2011).

Supporting this hypothesis, steep environmental gradients between upper gulf and open coastal environments corresponded with allele frequency differences at candidate adaptive loci. This included strong associations with temperature and salinity that are known to impact growth and survival of snapper (Fielder et al., 2005; McMahon et al., 2020). Previous snapper studies have demonstrated that adaptive traits such as growth and survival are strongly linked to these environmental variables (Wellenreuther et al., 2019), and have identified genomic variation underpinning these traits (Ashton et al., 2019a; Ashton et al., 2019b; Sandoval-Castillo et al., 2022). We found post-anal tail morphogenesis genes (*fgfr1a*, *tll1*, *bcl9l*) and haemopoietic regulators (*zfpm1*, *rnf145*, *smarcal1*) were significantly over-represented in our candidate adaptive loci. These functional modules align with growth-factor and oxygen-transport pathways that Wellenreuther et al. (2019) showed were sensitive to temperature in snapper, and have also been implicated in osmotic-stress responses in other fishes (Fiol & Kültz, 2007). This suggests that the same canonical signalling networks could mediate adaptation to the joint temperature-salinity gradients that span the embayment-coastal habitats. More broadly, the distribution of candidate adaptive loci across many regions of the genome is consistent with a signal of polygenic adaptation (Stephan, 2016).

While similar polygenic signals of selection have been observed in many marine species (Bernatchez, 2016), it is important to note that alternative genomic architectures have also been observed in highLgeneLflow marine fishes. In Atlantic silversides, strong divergent selection was shown to concentrate adaptive alleles into large genomic haploblocks (or “supergenes”) maintained by suppressed recombination, despite an overall low level of genomeLwide differentiation (Wilder, Palumbi, Conover, & Therkildsen, 2020). Structural variants can contribute to complex trait variation and can also create localised peaks of differentiation. Recent whole-genome analyses in snapper revealed substantial structural variation (Blommaert, Sandoval-Castillo, Beheregaray, & Wellenreuther, 2024) suggesting that future genomic studies may yield additional adaptive features beyond those our ddRAD approach could detect.

The role of natural selection in structuring marine populations across environmental gradients is gaining appreciation and this understanding is crucial for predicting climate-driven changes in fisheries sustainability (Teske et al., 2021). Local adaptation to temperature has been documented in economically important species such as the large yellow croaker (*Larimichthys crocea*), where genetic differentiation tracks minimum temperatures despite high connectivity, resulting in climate-induced shifts in stock boundaries (Chen et al., 2023). Similarly, salinity gradients were found to drive adaptive divergence in osmoregulation and reproductive traits among sand goby (*Pomatoschistus minutus*) populations across the North and Baltic Seas (Leder et al., 2021). Within Spencer Gulf and Gulf St. Vincent, where both temperature and salinity increase to the north, these environmental gradients influence multiple species. King George whiting (*Sillaginodes punctatus*), maintain genetic homogeneity across the South Australian region, yet comprise ecologically independent populations supplied by separate spawning grounds (Rogers, Fowler, Steer, & Gillanders, 2019). Despite their much longer pelagic larval phase (∼100 days, compared with 3-4 weeks in snapper), whiting exhibit the same fine-scale genetic structure linked to environmental variation at spawning grounds and to distinct larval dispersal routes. The recurrence of this pattern in taxa with contrasting life-history traits suggests a general pattern of environmental heterogeneity shaping fine-scale connectivity and population structure in marine fish populations across the region.

### Contingent migration can reinforce local adaptation with gene flow

Local adaptation across environmental gradients despite high gene flow has been documented in numerous marine species, including Atlantic cod (*Gadus morhua*), Pacific herring (*Clupea harengus*), and sardines (*Sardinops sagax*) (Bradbury et al., 2010; Limborg et al., 2012; Teske et al., 2021). These studies suggest that the balance between selection and gene flow in marine environments may be more nuanced than often assumed. Contingent migration, where populations contain both migratory and resident individuals, provides a potential general mechanism to explain the observed balance between connectivity and local adaptation (Hansson & Åkesson, 2014; Secor, 1999). Contingent migration strategies have been described for several marine fishes, including southern flounder (*Paralichthys lethostigma*) in the Gulf of Mexico (Steffen et al., 2023), and mulloway (*Argyrosomus japonicus*) and southern garfish (*Hyporhamphus melanochir*) in Australia (Hughes, Meadows, Stewart, Booth, & Fowler, 2022; Steer & Fowler, 2015). For snapper, tagging studies of the same population have revealed complex movement patterns with some individuals showing high site fidelity while others undertake extensive migrations (Fowler, 2016; Parsons et al., 2003; Stewart, Pidd, Fowler, & Sumpton, 2019). This is supported by evidence for three distinct behavioural groups in Port Phillip Bay snapper: fish that depart immediately after spawning, summer residents that remain until autumn, and individuals that overwinter in the bay (Hamer & Mills, 2017). Such behavioural variation is consistent with contingent migration where migratory individuals facilitate gene flow, while resident contingents maintain locally adapted genetic variation.

This balance between gene flow and selection is however unlikely to be temporally stable. The relative proportions of resident versus migratory contingents can shift in response to environmental variability and recruitment pulses, as documented in white perch (*Morone americana*) and striped bass (*M. saxatilis*) where contingent behaviour varied with river flow, productivity, and climatic conditions (Gahagan, Fox, & Secor, 2015; Gallagher, Piccoli, & Secor, 2018). For snapper, strong year classes might disperse more widely to reduce intra-specific competition, while weaker year classes may reinforce local residency (Drew et al., 2022). Previous genomic studies have detected temporal shifts in genetic composition, reflecting these variable recruitment and connectivity dynamics (Bertram et al., 2023).

Complex patterns of snapper stock structure observed in Shark Bay, Western Australia, further illustrate this temporal variation. Early tagging, otolith, and allozyme studies detected strong isolation between gulf and coastal populations (Jackson & Moran, 2012; Johnson et al., 1986), however subsequent microsatellite work revealed more subtle differentiation linked to salinity and timing of spawning (Gardner et al., 2017). These dynamics may also change under climate warming, potentially altering the timing and intensity of episodic recruitment or shifting the strength of environmental selection across heterogeneous spawning habitats. This highlights the importance of integrating knowledge about ecological and evolutionary processes into fisheries stock assessment and management, and into restocking and stock enhancement programs aimed at improving the abundance and resilience of wild stocks (e.g. Harrisson et al., 2025).

Our study provides novel insights into the complex relationship between gene flow and local adaptation in snapper populations along southern Australia’s environmentally heterogeneous coastline. The identification of genomic regions potentially under selection reveals restricted gene flow at candidate adaptive loci, highlighting that environmental factors shape population structure even in highly connected species like snapper. The potential role of contingent migration, where migratory and resident individuals coexist within populations, appears to be a key mechanism facilitating the balance between connectivity and local adaptation. This has direct implications for fisheries management in the region, particularly given the recent decline of some stocks. While current management necessarily operates within practical jurisdictional frameworks, our findings reveal biologically and ecologically relevant population structure that could enhance existing frameworks by incorporating both the demographic connectivity between regions and the adaptive distinctiveness of local populations. For instance, the Spencer Gulf and West Coast populations, currently managed as a single stock, show distinct adaptive signatures that may warrant consideration in refining future management strategies. Preserving these locally adapted populations that contribute to regional recruitment may prove crucial for persistence of snapper and other coastal species under climate change. While temporal variability in recruitment is already understood, longitudinal genomic monitoring could enhance assessment frameworks by revealing how this variability affects connectivity patterns and adaptive potential. Furthermore, strengthening collaboration between management jurisdictions would help ensure that population dynamics operating at large spatial scales are captured within local management strategies. As fisheries continue to experience unprecedented environmental changes, management strategies that integrate knowledge of both neutral and adaptive genetic variation, as well as of adaptive capacity, will be essential for ensuring long-term sustainability. Our study shows that demographic and adaptive connectivity can be decoupled in highly connected marine species and highlights that cryptic adaptive variation may prove crucial for population persistence in changing oceans.

## Acknowledgements

This study was financially supported by the Australian Research Council (grant LP180100756 to LBB), by Flinders University, and by the AJ and IM Naylon and the Playford Trust. We thank everyone who contributed towards sampling, including staff from the South Australian Research and Development Institute and the Victorian Fisheries Authority. We also thank Diana-Elena Vornicu for laboratory assistance. We acknowledge the traditional custodians of the land and sea throughout the regions where this research was done.

## Declaration of Interest Statement

The authors have nothing to declare.

## Data Accessibility and Benefit-Sharing

The study datasets are available in *Figshare*: https://figshare.com/s/034cb8ca31701cf9b687

Benefits from this research accrue from the sharing of our data and results on public databases as described above.

## Notes

https://figshare.com/s/034cb8ca31701cf9b687

### Competing Interest Statement

The authors have declared no competing interest.

