## Supplemental Files for "Environmental gradients decouple demographic and adaptive connectivity in a highly mobile coastal marine species"

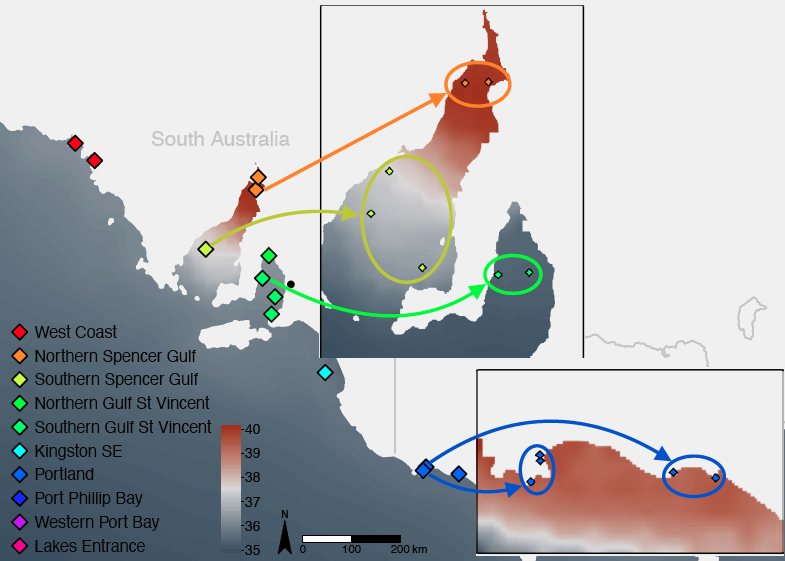

**Figure S1.** Sampling sites across the study area showing the 16 pooled sites (main panel) representing the 10 regional locations. Inset panels highlight individual sites that were pooled to ensure adequate sample sizes for allele-frequency estimation in a) northern Spencer Gulf, southern Spencer Gulf, southern Gulf St. Vincent and, b) Portland.

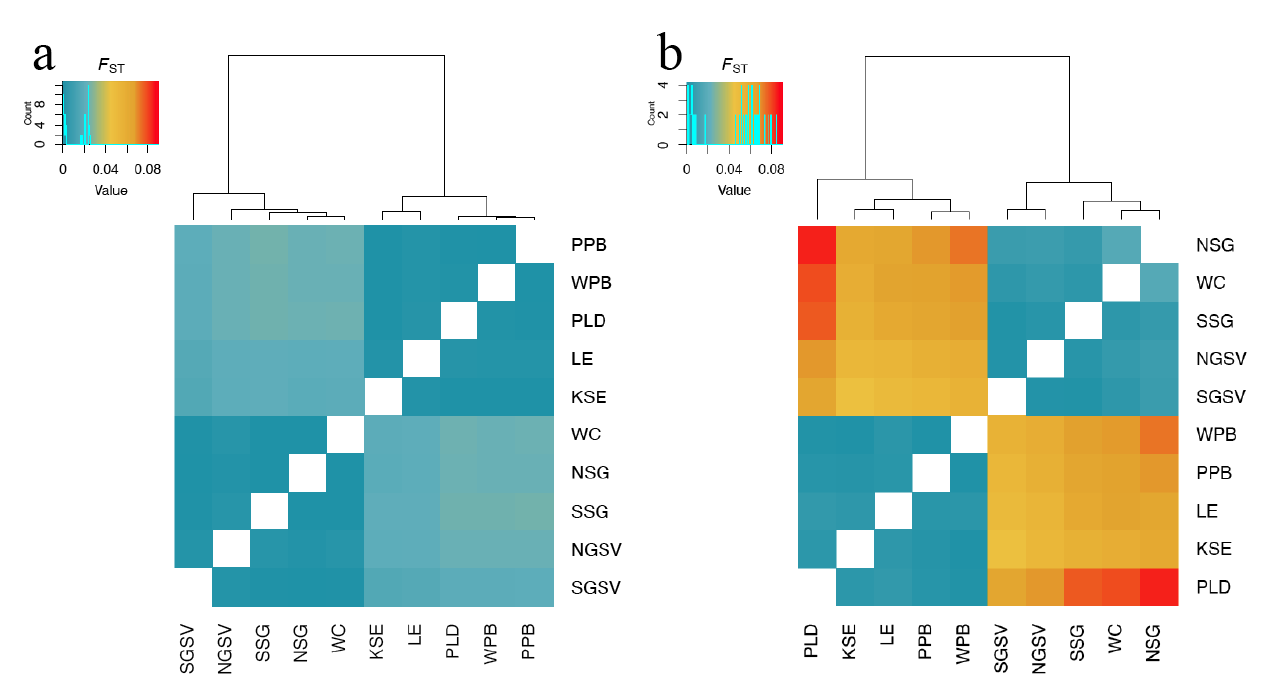

**Figure S2.** Pairwise *F*_ST_ based on (a) 13,453 neutral loci, and (b) 855 candidate adaptive loci.

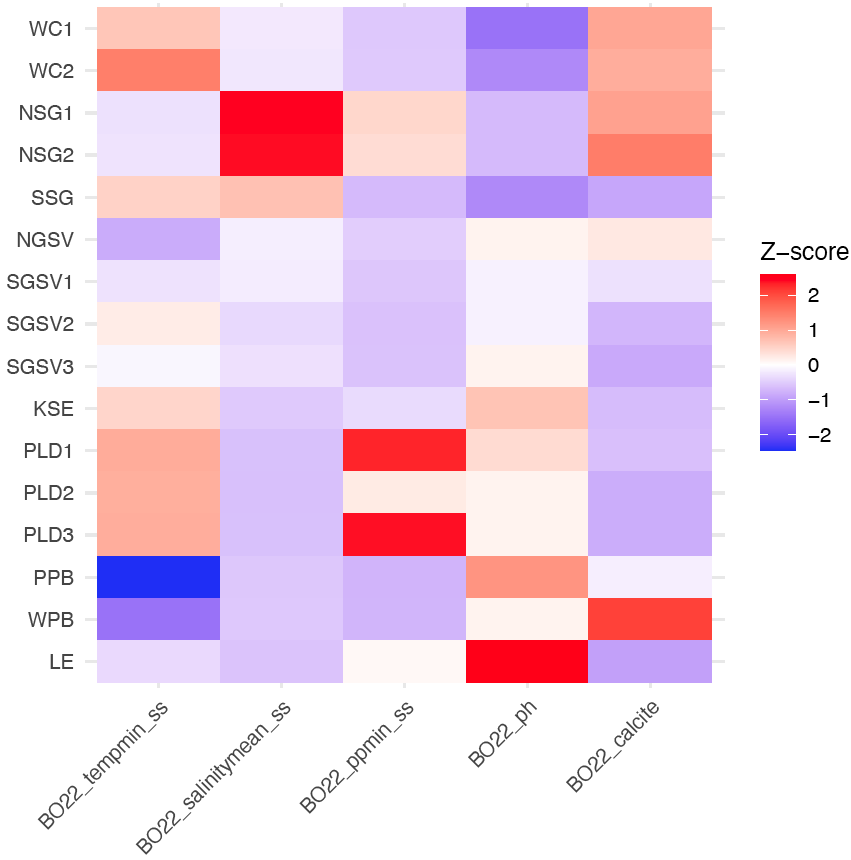

**Figure S3.** Environmental variation across the 16 pooled sites for snapper across southern Australia. Data are standardised with z-scores.

**Table S2.** Number of SNPs retained after each bioinformatics filtering step.

| Step | SNP count |
| --- | --- |
| Raw SNP catalogue | 81,810 |
| 80% of individuals, biallelic, >0.01 minor allele frequency | 72,766 |
| Mapping quality (>30) | 67,493 |
| Read quality (ratio quality/coverage depth >0.2) | 66,075 |
| High coverage loci (≤mean depth + (2*standard deviation)) | 66,055 |
| Linkage disequilibrium (500bp) | 14,699 |
| Hardy-Weinberg equilibrium (FDR 0.05) | 14,206 |
| Neutral loci | 13,453 |

**Table S4.** Relative migration estimates (Nm) for the eastern region based on 13,453 neutral loci.

|  | KSE | PLD | PPB | WPB | LE |
| --- | --- | --- | --- | --- | --- |
| KSE |  | 0.72 | 0.98 | 0.72 | 0.67 |
| PLD | 0.74 |  | 0.98 | 0.72 | 0.67 |
| PPB | 0.90 | 0.89 |  | 0.93 | 0.83 |
| WPB | 0.72 | 0.70 | 1.00 |  | 0.68 |
| LE | 0.68 | 0.66 | 0.88 | 0.69 |  |

**Table S5.** Relative migration estimates (Nm) for the western region based on 13,453 neutral loci.

|  | WC | NSG | SSG | NGSV | SGSV |
| --- | --- | --- | --- | --- | --- |
| WC |  | 0.74 | 0.76 | 0.60 | 0.92 |
| NSG | 0.77 |  | 0.76 | 0.63 | 0.97 |
| SSG | 0.76 | 0.73 |  | 0.64 | 1.00 |
| NGSV | 0.62 | 0.62 | 0.64 |  | 0.81 |
| SGSV | 0.87 | 0.87 | 0.95 | 0.76 |  |

**Table S6.** Relative migration estimates (Nm) for the eastern region based on 855 candidate loci.

|  | KSE | PLD | PPB | WPB | LE |
| --- | --- | --- | --- | --- | --- |
| KSE |  | 0.58 | 0.83 | 0.73 | 0.58 |
| PLD | 0.60 |  | 0.83 | 0.72 | 0.55 |
| PPB | 0.81 | 0.79 |  | 0.97 | 0.77 |
| WPB | 0.73 | 0.70 | 1.00 |  | 0.65 |
| LE | 0.58 | 0.54 | 0.80 | 0.64 |  |

**Table S7.** Relative migration estimates (Nm) for the western region based on 855 candidate loci.

|  | WC | NSG | SSG | NGSV | SGSV |
| --- | --- | --- | --- | --- | --- |
| WC |  | 0.32 | 0.62 | 0.50 | 0.68 |
| NSG | 0.33 |  | 0.52 | 0.45 | 0.54 |
| SSG | 0.63 | 0.51 |  | 0.68 | 1.00 |
| NGSV | 0.54 | 0.44 | 0.68 |  | 0.89 |
| SGSV | 0.69 | 0.51 | 0.96 | 0.83 |  |

**Table S8.** Raw environmental data for the 16 pooled snapper locations.

| Site | tempmin_ss | salinitymean_ss | ppmin_ss | ph | calcite |
| --- | --- | --- | --- | --- | --- |
| WC1 | 13.11 | 36.17 | 0.0004 | 8.251 | 0.0067 |
| WC2 | 13.74 | 36.16 | 0.0004 | 8.252 | 0.0064 |
| NSG1 | 12.37 | 40.05 | 0.0016 | 8.254 | 0.0069 |
| NSG2 | 12.39 | 39.97 | 0.0015 | 8.254 | 0.0081 |
| SSG | 13.00 | 37.51 | 0.0002 | 8.252 | 0.0009 |
| NGSV | 11.95 | 36.25 | 0.0004 | 8.257 | 0.0044 |
| SGSV1 | 12.39 | 36.22 | 0.0004 | 8.256 | 0.0027 |
| SGSV2 | 12.77 | 35.95 | 0.0003 | 8.256 | 0.0014 |
| SGSV3 | 12.53 | 36.07 | 0.0003 | 8.257 | 0.0010 |
| KSE | 12.97 | 35.72 | 0.0006 | 8.259 | 0.0016 |
| PLD1 | 13.34 | 35.59 | 0.0039 | 8.258 | 0.0017 |
| PLD2 | 13.32 | 35.59 | 0.0013 | 8.257 | 0.0011 |
| PLD3 | 13.33 | 35.60 | 0.0041 | 8.257 | 0.0011 |
| PPB | 10.70 | 35.69 | 0.0001 | 8.261 | 0.0031 |
| WPB | 11.44 | 35.71 | 0.0001 | 8.257 | 0.0100 |
| LE | 12.32 | 35.64 | 0.0012 | 8.266 | 0.0007 |
| Min | 10.70 | 35.59 | 0.0001 | 8.251 | 0.0007 |
| Max | 13.74 | 40.05 | 0.0041 | 8.266 | 0.0100 |
| Mean | 12.60 | 36.49 | 0.0011 | 8.256 | 0.0036 |
| sd | 0.78 | 1.45 | 0.0013 | 0.004 | 0.0030 |
